# Genetic Mapping Of Morpho-Physiological Traits Involved During Reproductive Stage Drought Tolerance In Rice

**DOI:** 10.1101/590075

**Authors:** Saumya Ranjan Barik, Elssa Pandit, Shakti Prakash Mohanty, Sharat Kumar Pradhan, Trilochan Mohapatra

**Author notes:** Corresponding author: Crop Improvement Division, ICAR-National Rice Research Institute, Cuttack, Odisha, India-753006.

## Abstract

Reproductive stage drought stress is an important factor for yield reduction in rice. Genetic mapping of drought responsive QTLs will help to develop cultivars suitable for drought prone environments through marker-assisted breeding. QTLs linked to morpho-physiological traits under drought stress were mapped by evaluating 190 F_7_ recombinant inbred lines (RIL). Significant variations were observed for eleven morpho-physiological traits involved during the stress. Bulked segregant analysis (BSA) strategy was adopted for genotyping the RIL population. A total of 401 SSR primers were tested for parental polymorphism of which 77 were polymorphic. Inclusive composite interval mapping detected a total of five consistent QTLs controlling leaf rolling (*qLR*_*9*.1_), leaf drying (*qLD*_*9*.1_), harvest index (*qHI*_*9*.1_), spikelet fertility (*qSF*_*9*.1_) and relative water content (*qRWC*_*9*.1_) under reproductive stage drought stress. Another two non-allelic QTLs controlling leaf rolling (*qLR*_*8*.1_) and leaf drying (*qLD*_*12*.1_) were linked in a single year. QTL controlling leaf rolling, *qLR8.1* was validated in this mapping population and useful in marker-assisted breeding (MAB) programs. Out of these five consistent QTLs, four (*qLR*_*9*.1_, *qLD*_*9*.1_, *qHI*_*9*.1_ and *qRWC*_*9*.1_) were detected to be novel QTLs and useful for MAB for reproductive stage drought tolerance in rice.

## Introduction

Majority of the global population consume rice as their staple food. Rice provides food and livelihood security to 90% of people in Asia and South-east Asia. Globally, rice crop covers about 160.8 million hectares with total production of more than 725.5 million tons of paddy per annum [1]. The targeted food production need to be increased even from the drought-prone areas with an increase of 40% from this difficult ecosystem by 2025 [2]. Drought is the major limitation in getting higher production from rainfed rice cultivation [3]. It affects the crop at vegetative and reproductive growth stages. Under water stress, genotype showing delay in leaf rolling and faster recovery from the stress are required for drought breeding [4]. During vegetative stage, leaf rolling and leaf drying are good criteria of screening drought tolerance [5–7]. Drought stress during reproductive stage is most critical as it causes low yield due to higher proportion of unfilled grains in the panicles [7, 9–12].

Drought is the major abiotic stress responsible for cultivation of rice mostly in rain-fed upland and lowland areas, worldwide. In Asia, about 34 million hectares of rain-fed lowland and 8 million hectare of upland rice are affected by frequent drought stress in each year. In India also, during the recent years, climate change is altering the rainfall pattern affecting rice production to a greater extent. Although rice production is increasing by 2.8% annually, damage caused by biotic and abiotic factors accounted for a heavy loss of global production during pre and post-harvest period [13].

Many primary, secondary and tertiary traits influence rice plant growth and production under drought stress. Majority of them are highly affected by reproductive stage drought stress. However, the information on genes/QTLs controlling various traits during the stress is not available or documented. A drastic decrease in grain yield and its component traits is usually seen due to the reproductive stage drought stress at a soil moisture level reaching to −60kPA or more. In addition to primary traits like grain yield and root parameters, many morphological and physiological traits and production change are associated with the stress level. Reproductive stage is regarded as the most crucial stage of drought stress in compare to vegetative stage drought [8,14]. The responses to drought in the vegetative stage and reproductive stage are much different [15]. Several experiments have been reported previously for morpho-physiological responses under reproductive stage drought stress [16–27].

Though sporadic research works reported on few QTLs linked to reproductive stage drought tolerance, but no robust markers controlling traits under this stress are available. CR 143-2-2, a drought donor line possessing both vegetative and reproductive stage drought tolerance and the contrasting parent Krishnahamsa (susceptible variety for drought) were used for development of mapping population. As reproductive stage drought stress is responsible for drastic reduction in yield, mapping of QTLs responsive to the stress will help to develop rice cultivars suitable for drought prone environments through marker-assisted breeding. Therefore, QTLs linked to morpho-physiological traits under drought stress were mapped by phenotyping 190 F_7_ recombinant inbred lines (RIL) with the polymorphic markers using bulk segregant analysis.

## Materials and methods

### Plant materials

One hundred ninety recombinant inbred lines (F_7_:F_8_) were selected and maintained from the cross between tolerant parent CR 143-2-2 and agronomic important parent Krishnahamsa. Pure line selection by single seed decent (SSD) method was followed in each generation (F_2_-F_8_) in normal field condition. Experiments for recombinant lines were carried out under controlled facilities of Rain-out shelter (RoS) in ICAR-National Rice Research Institute (NRRI) in two consecutive years i.e. wet season, 2014 and 2015. Developed recombinant lines were subjected to phenotyping and genotyping study at F_7-8_ generation. CR 143-2-2 is the donor parent developed by ICAR-National Rice Research Institute for upland ecology breeding system where as Krishnahamsa (DRR Dhan 20) is the agronomical important parent developed by Indian Institute of Rice Research, Hyderabad which is a high yielding popular irrigated variety of Andhra Pradesh state of India.

### Phenotyping for physiological traits under reproductive stage drought stress

The field experiment was conducted by direct sowing the seeds of recombinant lines in alpha lattice design with two replications in rain out shelter. The seeds were shown uniformly in August month of each year and maintained in normal condition by irrigation up to completion of vegetative stage. All the recombinant lines along with their parents were arranged in six blocks constituting 34 lines per block with spacing of 10X15 cm (row per hill). Each row consisting of 25 hills per each recombinant lines from which 10 hill sample data were taken up for the evaluation process. Drought stress was applied at the very early PI (primordium initiation) stage simultaneously to all the recombinant lines and the parents. Doses for fertilizer application were maintained in appropriate time interval with a ratio of 40:10:10 kg/ha for nitrogen (N): phosphorous (P): potassium (K) respectively. Throughout the reproductive stage, drought stress was maintained continuously up to −50kPA in the experimental field condition. Eleven physiological traits *viz*., plant height, leaf rolling, leaf drying, panicle length, percentage of panicle emergence, harvest index, 1000-grain weight, percentage of spikelet fertility, relative water content, cell membrane stability and grain yield were estimated under the stress condition.

Data for plant height of each recombinant lines and parents were recorded at growth stage 7. Other pre-harvested data *viz*., leaf rolling, leaf drying, panicle length and percentage of panicle emergence were collected during growth stage of 6-9 [28]. From 10 hills sample collection, post-harvest data *viz*., harvest index, thousand grain weight, percentage of spikelet fertility and grain yield were collected recorded at growth stage 9. For estimation of physiological traits, samples for relative water content and cell membrane stability were collected from the field at mid-day situation at growth stage 7-8. Estimation of relative water content was done as per the method of [29]. Samples for cell membrane stability were collected during mid-day period following the procedure of [30].

### Genotyping

#### DNA extraction

Fresh young leaf samples (20-25 days) were collected from the population aseptically for genomic DNA extraction. Leaves were homogenized with the help of liquid nitrogen in mortar and pistle collected in 2ml micro centrifugation tube. Pre warmed (65°C) Cetyltrimethyl ammonium bromide (CTAB) extraction buffer (2% CTAB, 100mM Tris pH 8, 20mM Ethylene diamine tetra acetate (EDTA) pH 8, 1.3M NaCl) was added to the sample followed by phenol chloroform isoamyl alcohol (PCI) extraction, RNase treatment and ethanol precipitation [31]. The quality and quantity of final extracted DNA was verified with λ-DNA on 1% agarose gel. Also, DNA quantification and purity was further checked by measuring the OD at 260 and 280nm using a UV visible spectrophotometer. The samples were diluted accordingly for their uniformity to approximately 30ng/μl.

#### Polymerase chain reaction (PCR)

The polymerase chain reaction was performed by taking 20μl aliquot in a programmable thermal cycler (Veriti, Applied-Biosystems) using SSR primers (Table 4; Table 5). The PCR reaction mixture included 30ng genomic DNA, 10mM dNTPs, 2mM MgCl_2_, 50mM KCL, 1.5 mM Tris HCL (pH 8.75), 1U Taq polymerase and 10μM each of forward and reverse primers. The thermal profile starts with initial denaturation at 94°C for 4 min continued to 35 cycles of denaturation at 94°C for 30 sec, primer annealing 55°C for 1 min, extension 72°C for 1.30 min and final extension at 72°C for 10 min. After completion of amplification, PCR products were kept at −20°C. 10 μl aliquot of PCR amplification products were load in an agarose gel of 3.5% containing 0.08 μg/ml of ethidium bromide for electrophoresis. Sizes of amplicon were determined by using 50bp DNA ladder. The electrophoresis was carried out at 80 volts (2.5 V/cm) in 1X TBE (pH 8.0). The photograph of banding pattern was documented using gel documentation unit (Syngene G Box).

#### Bulked segregant analysis

The method for bulked segregant analysis (BSA) is used to detect the major QTL linked to the trait of interest [32]. From the experimental result of phenotypic classification for all 190 recombinant inbred line populations, ten recombinant lines were selected from each extreme tolerant and susceptible gene pools and bulked to detect the molecular variation using polymorphic SSR markers. Detected polymorphic markers through BSA analysis and their relation with phenotypic effects were further analyzed through molecular mapping method using ICIM v4.0 software.

#### Statistical analysis

Analysis of mean, range, skewness and kurtosis of all 190 recombinant lines and their parents were estimated from the two year data *i.e. kharif*, 2014 and 2015. These phenotypic distributions among the recombinant lines for relative traits to determining the main effect was performed by using software SPSS v20.0 [33]. INDOSTAT software [34] was used to determine the correlation analysis and genetic advance among the recombinant lines. Coefficient of variance (CV) and LSD_5%_ were obtained by using the software CROPSTAT v7.0. Also, phenotypic data analysis like phenotypic covariance (PC), genotypic covariance (GC), environmental covariance (EC), phenotypic coefficient of variance (PCV), genotypic coefficient of variance (GCV) and heritability (H^2^) value were performed by the software SPAR v2.0.

#### Linkage map and QTL analysis

Phenotypic data of eleven traits *viz*., plant height, leaf rolling, leaf drying, panicle length, percentage of panicle emergence, harvest index, thousand seed weight, percentage of spikelet fertility, relative water content, cell membrane stability and grain yield along with genotypic data of 190 recombinant lines and parents were utilize for construction of linkage map by using the software inclusive composite interval mapping v4.0 (ICIM v4.0) [33]. Analysis of composite interval mapping (CIM) and additive effect in relation to QTL mapping were used to calculate the association of phenotypic and molecular proportions for the development of linkage map. The walking speed along chromosomes for all QTLs was 1.0cM, and threshold value of LOD 2.0 with 1000 permutation for P<0.05 were considered. The QTLs were named according to the nomenclatural guidelines given in [35].

## Results

### Variation in morpho-physiological traits of RILs under reproductive stage drought stress

Drought stress affects growth and development of rice plant particularly at flowering stage thereby reduces yield. During wet seasons 2014 and 2015, eleven morpho-physiological traits were estimated under reproductive stage drought stress. Significant variations were obtained from the contrasting parents for all the 11 mopho-physiological traits (Table 1). All the morpho-physiological parameters estimates *viz*., percentage of panicle emergence, harvest index, 1000-grain weight, percentage of spikelet fertility, relative water content, cell membrane stability and grain yield were observed to be high in tolerant parent, CR 143-2-2 than susceptible parent Krishnahamsa except plant height, leaf rolling, leaf drying and panicle length. Thus, the selection of contrasting parents for development of mapping population for mapping of these traits were effective.

**Table 1.** Mean estimates of morpho-physiological traits in contrasting parents (CR143-2-2 and Krishnahamsa) under reproductive stage drought stress

| Sl. No. | Phenotyping traits | CR 143-2-2<br>(Tolerant Parent) | Krishnahamsa<br>(Susceptible Parent) |
| --- | --- | --- | --- |
| 1 | Plant height | 66cm | 74cm |
| 2 | Leaf rolling (SES score) | 0.5 | 6.0 |
| 3 | Leaf drying (SES score) | 0.5 | 5.5 |
| 4 | Panicle length | 18.81cm | 20.94cm |
| 5 | Panicle emergence (%) | 95% | 93% |
| 6 | Harvest index | 0.48 | 0.234 |
| 7 | 1000- seed weight (g) | 22.94 | 20.25 |
| 8 | Spikelet fertility (%) | 84.62 | 55.10 |
| 9 | Relative water content (%) | 90.57 | 56.78 |
| 10 | Cell membrane stability (%) | 86.73 | 51.14 |
| 11 | Grain yield (g) | 5.82 | 1.23 |

Plant height (PH) is an important parameter controlling plant type of RILs in stress condition. In our study, significant variation was obtained among the recombinant lines for this trait (Table 2). In our experiment plant height shows a larger variation which ranged from 45.67cm to 117.92cm. Coefficient of variations in the RILs obtained for PH, 8.3 was quite significant (Table 2). While CR 143-2-2 has shorter plant height of 66cm in contrast to Krishnahamsa which has longer height of 74cm (Table 1).

**Table 2.** Mean statistical parameters for eleven morpho-physiological traits under reproductive stage drought stress

| Trait | Mean | Range | Skewness | Kurtosis | CV (%) | LSD<sub>(5%)</sub> | PCV (%) | GCV (%) | H<sup>2</sup> (%) |
| --- | --- | --- | --- | --- | --- | --- | --- | --- | --- |
| **PH (cm)** | 83.96 | 45.67-117.92 | 0.008 | 0.16 | 6.3 | 13.79 | 16.16 | 13.84 | 0.73 |
| **LR** | 2.8 | 0-9.0 | 0.92 | 0.12 | 12.5 | 0.95 | 76.34 | 72.93 | 0.91 |
| **LD** | 1.57 | 0-7.0 | 1.80 | 2.43 | 14.1 | 0.62 | 108.02 | 103.44 | 0.92 |
| **PL (cm)** | 20.12 | 14.38-25.75 | 0.014 | 0.005 | 9.8 | 3.89 | 11.84 | 7.64 | 0.65 |
| **PE (%)** | 95.63 | 80-100 | -1.25 | 3.55 | 3.7 | 6.98 | 4.20 | 1.97 | 0.22 |
| **HI (%)** | 0.4 | 0.152-0.583 | -0.23 | 0.71 | 15.4 | 0.08 | 21.77 | 18.43 | 0.85 |
| **TSW(g)** | 20.25 | 5.51-33.22 | -0.8 | 1.98 | 6.0 | 2.38 | 21.14 | 20.28 | 0.92 |
| **SF (%)** | 74.02 | 44.68-91.96 | -0.78 | 0.24 | 16.8 | 18.48 | 18.03 | 12.64 | 0.70 |
| **RWC (%)** | 75.52 | 12.08-98.16 | 0.18 | 1.18 | 5.9 | 8.74 | 21.45 | 20.63 | 0.93 |
| **CMS** | 63.42 | 4.29-91.52 | -0.78 | 0.4 | 10.0 | 12.54 | 28.59 | 26.78 | 0.88 |
| **YLD (g)** | 3.75 | 0.82-7.68 | 0.79 | 1.38 | 17 | 0.92 | 44.22 | 33.46 | 0.76 |

PH=plant height, LR= leaf rolling, LD= leaf drying, PL= panicle length, PE= % of panicle emergence, HI= harvest index, TSW=thousand seed weight, SF= spikelet fertility, RWC= relative water content, CMS= cell membrane stability, YLD= grain yield, CV= coefficient of variation, LSD= least square deviation, PCV= phenotypic coefficient of variation, GCV= genotypic coefficient of variation, H<sup>2</sup>= heritability (broad sense)

Leaf rolling (LR) and leaf drying (LD) are the most important traits for identifying drought donor parents. The tolerant donor, CR 143-2-2 showed very low SES score compared to the susceptible parent, Krishnahamsa (Table 1). In this investigation, a wide variation in LR was observed showing a range of 0 to 9.0 with mean of 2.8 among the RILs. Similarly, LD also showed a wide variation with mean value of 1.57 among the RILs under drought stress condition (Table 2). Coefficient of variation and LSD_5%_ of LR and LD were observed to be 12.5, 0.95 and 14.1, 2, respectively. A very high heritability of 91 and 92% were computed for both LR and LD, respectively.

The traits, panicle length (PL) and panicle emergence (PE) showed variations in both the parents and among RILs. Panicle length had a range of 14.38to 25.75cm, whereas panicle emergence showed 80-100% among RILs. Under reproductive stage drought stress, both the parents showed a clear variation for the trait harvest index (HI) and grain yield (YLD). CR 143-2-2 showed higher values of 0.48 (HI value) while 0.234 (HI value) was in susceptible parent, Krishnahamsa. Single plant yield (YLD) obtained from CR 143-2-2 (5.82g) was much higher compared to Krishnahamsa (1.23gm). HI showed a wide range of 0.152 to 0.583 with a mean of 0.4 whereas YLD ranged from 0.82 to 5.68 with mean of 3.75 for all the RILs under drought. Coefficient of variation and LSD_5%_ for HI and YLD values of 15.4, 0.12 and 22.5, 1.22, respectively. A higher heritability (broad sense) was also computed for both the traits.

A wide variation in 1000-seed weight (TSW), spikelet fertility (SF) and relative water content (RWC) was observed in both the parents, CR143-2-2 and Krishnahamsa (Table 1). The range of TSW, SF and RWC in RILs also exhibited a variation of 5.51-33.22, 44.68-91.96 and 12.08-98.16 with a mean of 20.25, 74.02 and 75.52, respectively. Heritability (broad sense) values were observed for both the traits, TSW (92%) and RWC (93%). The cell membrane stability (CMS) showed a range of 6.74-93.81 with a mean value of 63.42 for the RILs. The donor parent, CR 143-2-2 showed a higher estimate of CMS value compared to susceptible parent, Krishnahamsa. Also, the heritability (broad-sense) was high for the trait (Table 2).

### Frequency distributions

The frequency distributions of 190 RILs along with parents for all the eleven morpho-physiological traits are depicted in the Fig. 1. Wide variation was observed for each trait in the parental lines depicted as P1 (tolerant parent) and P2 (susceptible parent). Skewness and kurtosis values of the respective identified traits for a normal curve are provided in Table 2. With both the positive values of skewness and kurtosis, six morpho-physiological traits *viz*., plant height, leaf rolling, leaf drying, panicle length, relative water content and grain yield showed a positive leptokurtic skewed distribution. However, rest of five traits *viz*., panicle emergence, harvest index, 1000-seed weight, spikelet fertility and cell membrane stability exhibited a positive kurtosis value and negative skewness value showing negatively skewed leptokurtic distribution. Under drought condition, behavioral pattern of plant might be changed due to stress effect which was seen in the distribution study of leaf drying, leaf rolling and panicle emergence traits. Rest of the eight traits showed almost a normal distribution curve (Fig. 1).

**Fig 1.**
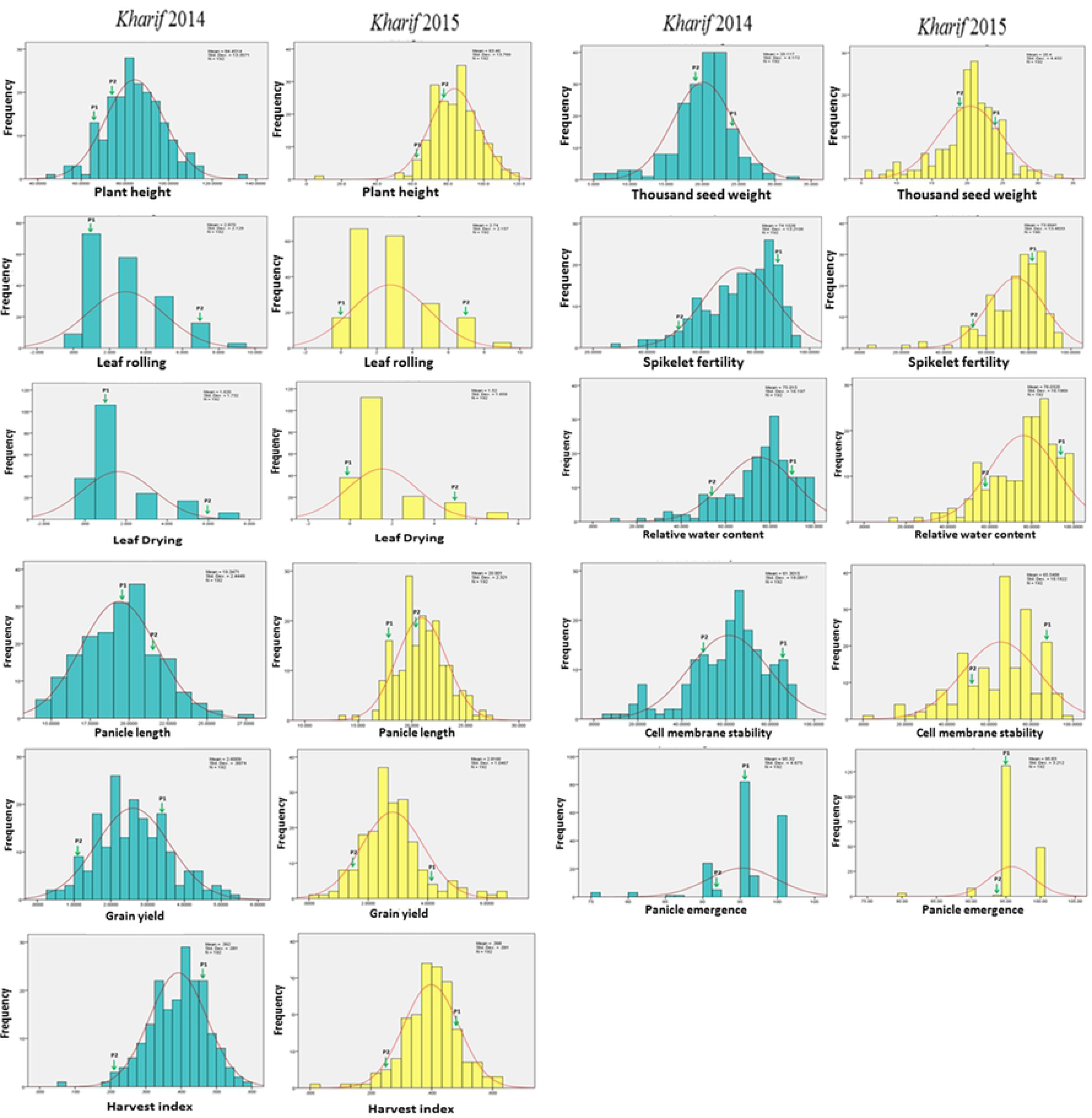
Frequency histogram and phenotypic normal distribution curves of all eleven phenotypic traits

### Correlation of morpho-physiological traits with grain yield under reproductive stage drought stress

The correlation coefficients of eleven studied morpho-physiological traits among themselves and with grain yield were observed to be significant under the stress (Table 3). Out of the total 55 correlations, 25 correlations were significant at the level of 0.01 and 5 correlations were significant at the level of 0.05. A very positive high correlation value was observed for HI and grain yield (r=0.865**) followed by –ve significant correlation of leaf rolling and leaf (Table 3). Also, a strong negative correlation was observed between leaf drying and harvest index (r = −0.203**) at 0.01 level of probability. Under stress condition, grain yield showed significant positive correlation with panicle length, plant height, panicle emergence, harvest index, 1000-seed weight, spikelet fertility and relative water content (Table 3).

**Table 3.**
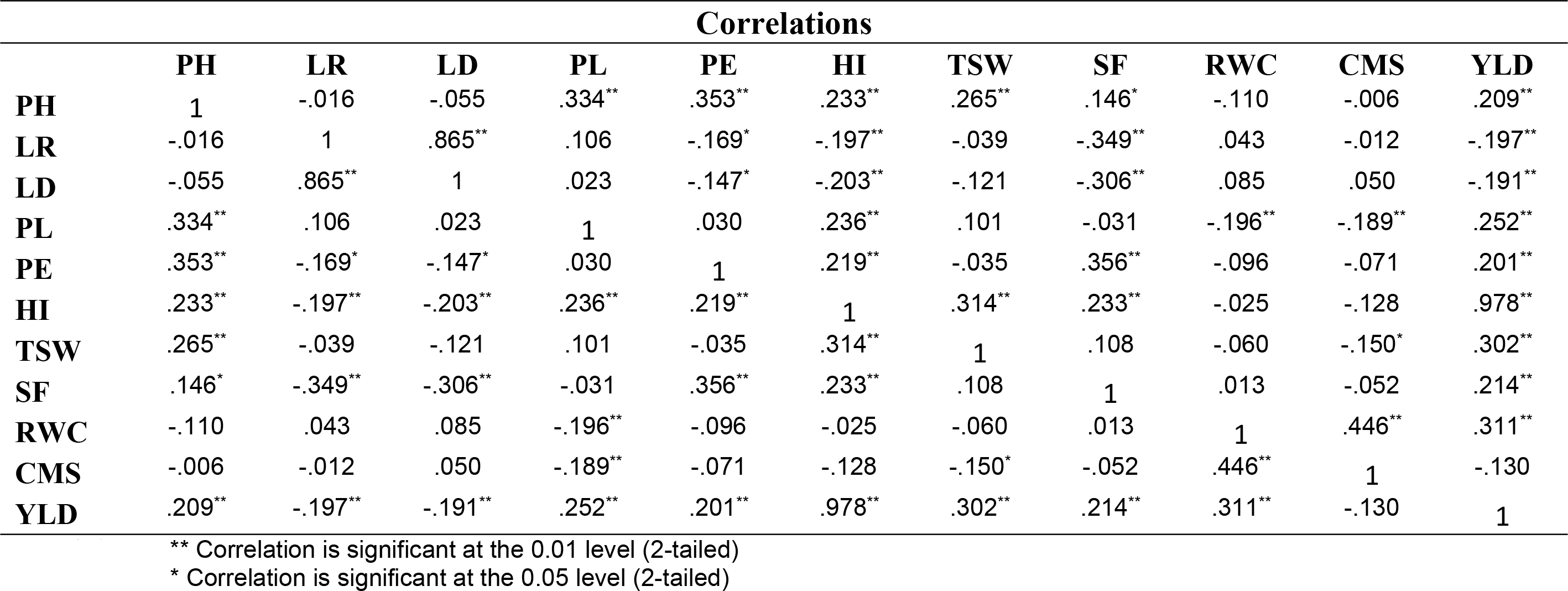
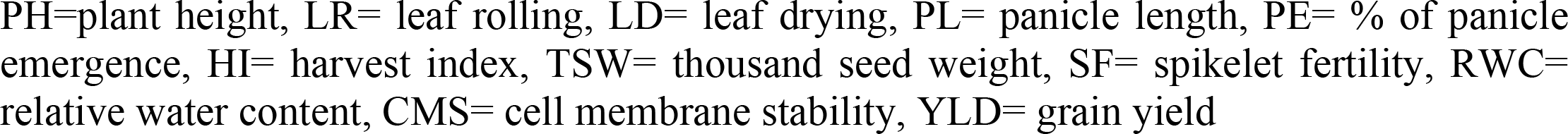
Correlation of morpho-physiological traits with grain yield under reproductive drought stress

| Correlations |  |  |  |  |  |  |  |  |  |  |  |
| --- | --- | --- | --- | --- | --- | --- | --- | --- | --- | --- | --- |
|  | PH | LR | LD | PL | PE | HI | TSW | SF | RWC | CMS | YLD |
| **PH** | 1 | -.016 | -.055 | .334** | .353** | .233** | .265** | .146* | -.110 | -.006 | .209** |
| **LR** | -.016 | 1 | .865** | .106 | -.169* | -.197** | -.039 | -.349** | .043 | -.012 | -.197** |
| **LD** | -.055 | .865** | 1 | .023 | -.147* | -.203** | -.121 | -.306** | .085 | .050 | -.191** |
| **PL** | .334** | .106 | .023 | 1 | .030 | .236** | .101 | -.031 | -.196** | -.189** | .252** |
| **PE** | .353** | -.169* | -.147* | .030 | 1 | .219** | -.035 | .356** | -.096 | -.071 | .201** |
| **HI** | .233** | -.197** | -.203** | .236** | .219** | 1 | .314** | .233** | -.025 | -.128 | .978** |
| **TSW** | .265** | -.039 | -.121 | .101 | -.035 | .314** | 1 | .108 | -.060 | -.150* | .302** |
| **SF** | .146* | -.349** | -.306** | -.031 | .356** | .233** | .108 | 1 | .013 | -.052 | .214** |
| **RWC** | -.110 | .043 | .085 | -.196** | -.096 | -.025 | -.060 | .013 | 1 | .446** | .311** |
| **CMS** | -.006 | -.012 | .050 | -.189** | -.071 | -.128 | -.150* | -.052 | .446** | 1 | -.130 |
| **YLD** | .209** | -.197** | -.191** | .252** | .201** | .978** | .302** | .214** | .311** | -.130 | 1 |

\*\* Correlation is significant at the 0.01 level (2-tailed)
\* Correlation is significant at the 0.05 level (2-tailed)
PH=plant height, LR= leaf rolling, LD= leaf drying, PL= panicle length, PE= % of panicle emergence, HI= harvest index, TSW= thousand seed weight, SF= spikelet fertility, RWC= relative water content, CMS= cell membrane stability, YLD= grain yield

### QTL mapping of morpho-physiological traits under reproductive stage drought stress

In this present investigation, 401 SSR primers were utilized for detection of parental polymorphism (Table 4). Out of 401 primers, 77 primers were polymorphic in the parents. For bulked segregant analysis (BSA), two different extreme RILs bulks (tolerant: B1 and susceptible: B2) were constructed and utilized for genotyping with the 77 polymorphic primers (Table 5; Fig. 2).

**Table 4.** Microsatellite markers obtained through the polymorphic analysis between CR143-2-2 and Krishnahamsa

| Chromosome | No. of markers analyzed | Total No. and names of the parental polymorphic markers obtained |  | Total No. and names of the bulked polymorphic markers used |  |
| --- | --- | --- | --- | --- | --- |
| 1 | 50 | 10 | RM6703, RM3825, RM488, RM259, RM5, RM12091, RM8085, RM495, RM5443, RM1003 | 3 | RM495, RM6703, RM3825 |
| 2 | 50 | 11 | RM324, RM263, RM327, RM530, RM262, RM3549, RM279, OSR17, RM250, RM 341, RM13600 | 3 | RM327, RM341, RM263 |
| 3 | 42 | 12 | RM523, RM231, RM7332, RM517, RM411, RM135, RM85, RM22, RM16030, RM15780, RM104, RM571 | 2 | RM22, RM517 |
| 4 | 26 | -- | -- | -- | -- |
| 5 | 8 | -- | -- | -- | -- |
| 6 | 30 | 4 | RM3, RM276, RM527, RM528 | 2 | RM527, RM3 |
| 7 | 14 | 1 | MGR4499 | -- | -- |
| 8 | 22 | 6 | RM256, RM337, RM210, RM25, RM342A, RM 72 | 2 | RM337, RM72 |
| 9 | 38 | 7 | RM464, RM215, RM219, RM316, RM257, RM242, RM213 | 2 | RM316, RM257 |
| 10 | 26 | 6 | RM216, RM228, RM311, RM271, RM171, RM484 | 3 | RM271, RM171, RM484 |
| 11 | 10 | 1 | RM21 | -- | -- |
| 12 | 84 | 19 | RM28199, RM28089, RM511, RM28166, RM1261, RM28048, RM28059, RM28064, RM28067, RM28070, RM28079, RM28082, RM28083, RM28088, RM28090, RM519, RM313, RM309, RM20A | 4 | RM20A, RM511, RM309, RM519 |
| Total | 401 | 77 |  | 21 |  |

**Table 5.**
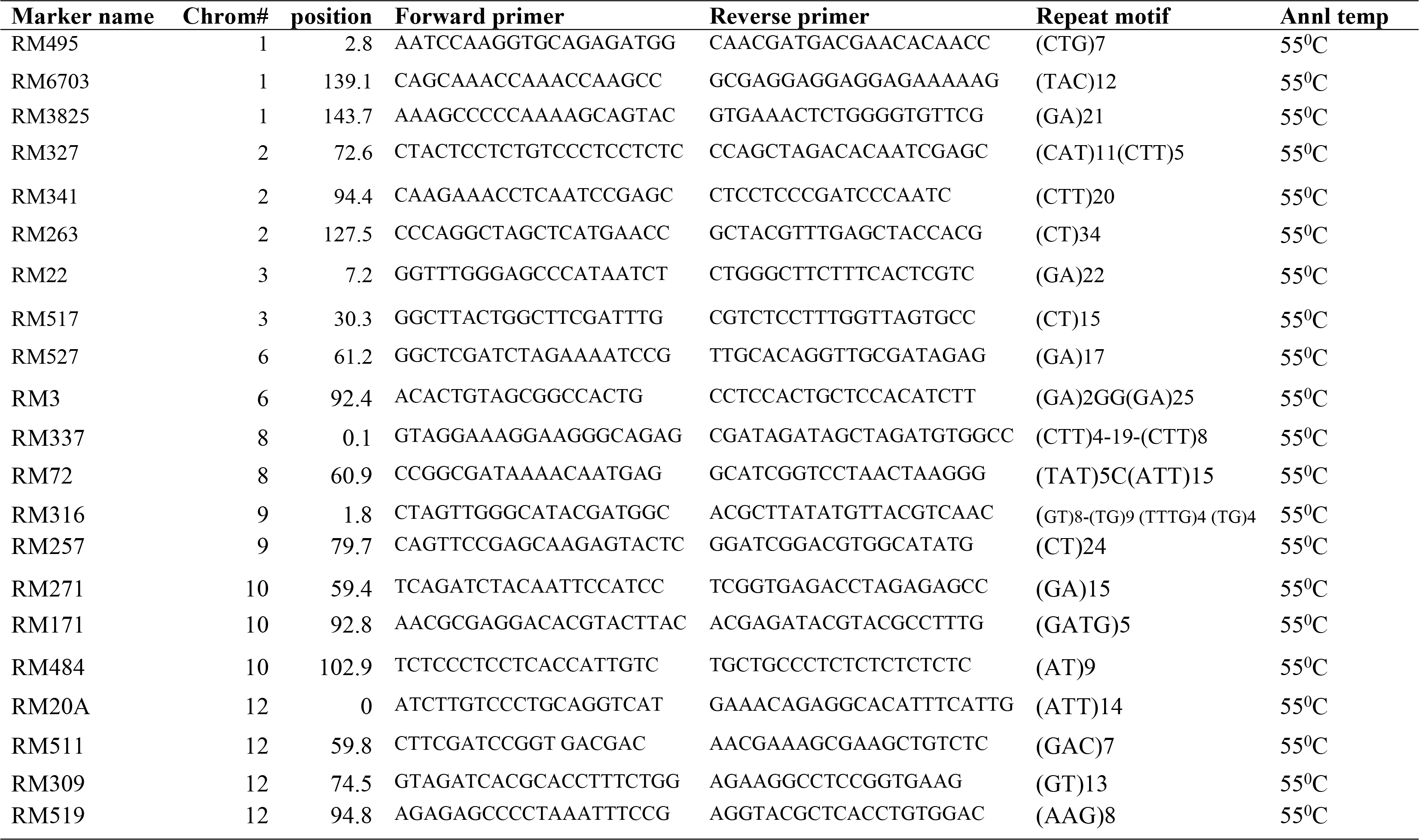
Details of polymorphic SSR markers detected in the RILs population

**Fig 2.**
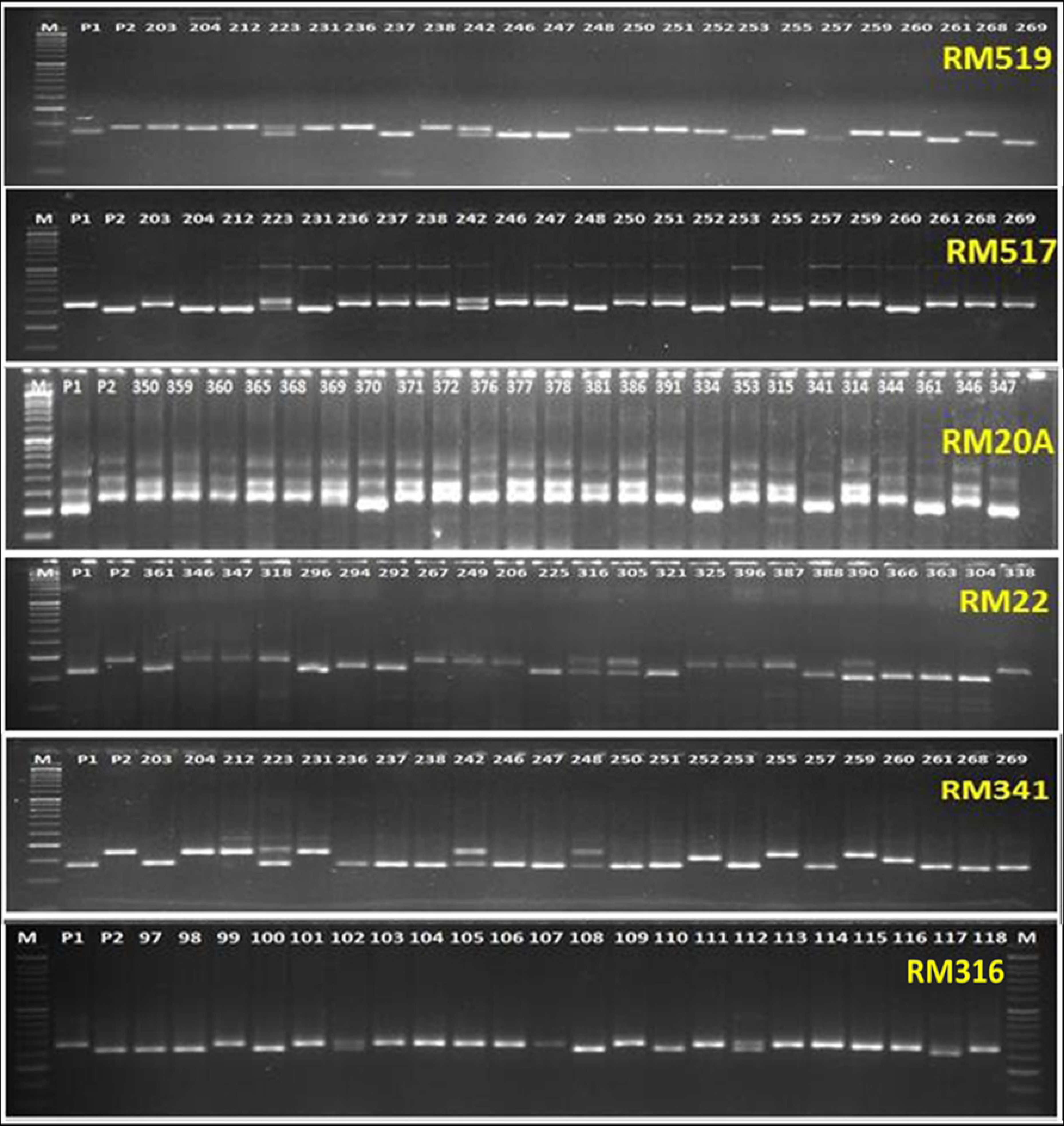
Electrophoregram showing polymorphism pattern of SSR primers with different recombinant lines. The numbers represent the different RI line numbers used in mapping. Respective primer names are given in the right top corner position in each gel photos. **P1:** Tolerant parent; **P2:** Susceptible parent, **M:** 50bp DNA ladder

The analysis using inclusive composite interval mapping (ICIM) method detected seven QTLs linked to leaf drying, leaf rolling, harvest index, spikelet fertility and relative water content traits on three different chromosomes (Chromosome 8, 9 and 12) in year I (wet season, 2014),. However, in year II (wet season, 2015), five QTLs found to be linked to same traits in one chromosomes (Chromosome 9) under drought stress condition (Table 6; Fig. 3B). These seven QTLs (LOD ≥ 2.5) represented five different morpho-physiological traits and observed to be located in three different chromosomes.

**Table 6.** QTL identified from inclusive composite interval mapping

| Year | QTL detected | Chrom# | Position | LM | RM | LOD | PVE(%) | Additive effect |
| --- | --- | --- | --- | --- | --- | --- | --- | --- |
| Kharif,2014 | qLR<sub>8.1</sub> | 8 | 27.1 | RM337 | RM72 | 2.78 | 60.74 | -1.68 |
| Kharif,2015 | - | - | - | - | - | - | - | - |
| Pooled | - | - | - | - | - | - | - | - |
| Kharif,2014 | qLR<sub>9.1</sub> | 9 | 25.8 | RM316 | RM257 | 7.20 | 67.46 | -1.90 |
| Kharif,2015 | qLR<sub>9.1</sub> | 9 | 19.8 | RM316 | RM257 | 5.41 | 59.03 | -1.80 |
| Pooled | qLR<sub>9.1</sub> | 9 | 22.8 | RM316 | RM257 | 8.21 | 65.99 | -1.82 |
| Kharif,2014 | qLD<sub>9.1</sub> | 9 | 13.8 | RM316 | RM257 | 17.18 | 70.34 | -1.94 |
| Kharif,2015 | qLD<sub>9.1</sub> | 9 | 15.8 | RM316 | RM257 | 21.85 | 77.86 | -1.64 |
| Pooled | qLD<sub>9.1</sub> | 9 | 14.8 | RM316 | RM257 | 27.85 | 80.18 | -1.81 |
| Kharif,2014 | qLD<sub>12.1</sub> | 12 | 45.0 | RM20A | RM511 | 11.35 | 71.72 | 1.82 |
| Kharif,2015 | - | - | - | - | - | - | - | - |
| Pooled | - | - | - | - | - | - | - | - |
| Kharif,2014 | qHI<sub>9.1</sub> | 9 | 12.8 | RM316 | RM257 | 3.51 | 52.40 | 0.01 |
| Kharif,2015 | qHI<sub>9.1</sub> | 9 | 12.8 | RM316 | RM257 | 7.02 | 60.20 | 0.01 |
| Pooled | qHI<sub>9.1</sub> | 9 | 11.8 | RM316 | RM257 | 5.21 | 56.45 | 0.01 |
| Kharif,2014 | qSF<sub>9.1</sub> | 9 | 35.8 | RM316 | RM257 | 2.26 | 49.85 | 9.88 |
| Kharif,2015 | qSF<sub>9.1</sub> | 9 | 12.8 | RM316 | RM257 | 3.50 | 45.79 | 11.68 |
| Pooled | qSF<sub>9.1</sub> | 9 | 23.8 | RM316 | RM257 | 3.58 | 54.5 | 8.35 |
| Kharif,2014 | qRWC<sub>9.1</sub> | 9 | 18.8 | RM316 | RM257 | 4.13 | 59.62 | 15.14 |
| Kharif,2015 | qRWC<sub>9.1</sub> | 9 | 23.8 | RM316 | RM257 | 4.78 | 62.65 | 14.52 |
| Pooled | qRWC<sub>9.1</sub> | 9 | 21.8 | RM316 | RM257 | 4.27 | 60.87 | 14.56 |

LR= leaf rolling, LD= leaf drying, HI= harvest index, SF= spikelet fertility, RWC= relative water content, LM= left marker, RM= right marker

**Fig 3.**
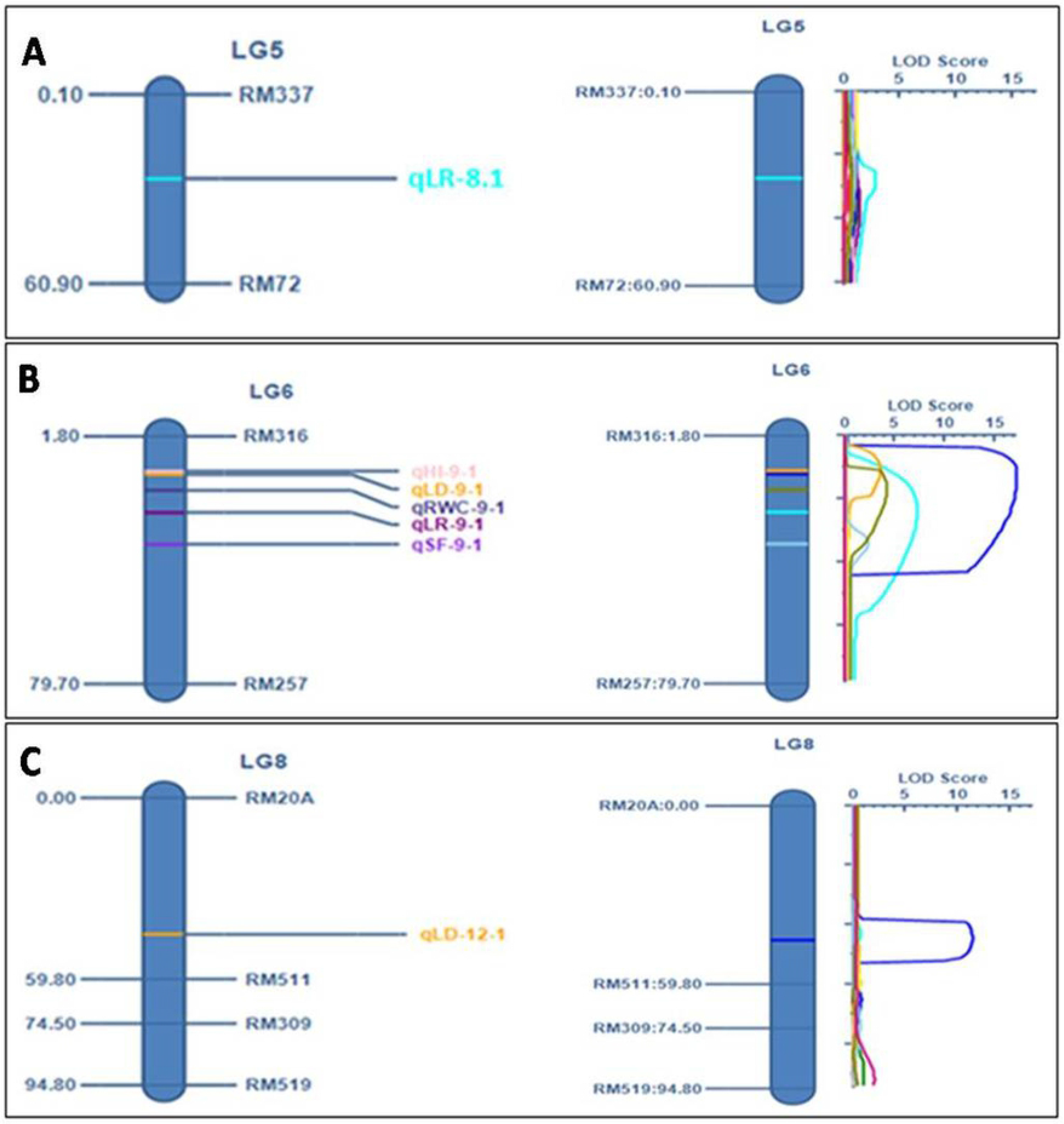
(A) QTL detected on chromosome 8 (LG5) beyond threshold LOD (2.5) (light green color represents QTL detected for leaf rolling) (B) QTL detected on chromosome 9 (LG6) beyond threshold LOD (2.5) (pink color represents QTL detected for harvest Index, yellow color represents QTL detected for leaf drying, deep blue represents QTL detected for relative water content, maroon color represents QTL detected for leaf rolling and violet color represents QTL detected for spikelet fertility) (C) QTL detected on chromosome 12 (LG8) beyond threshold LOD (2.5) (yellow color represents QTL detected for leaf drying)

The QTL for leaf rolling was detected on both chromosome 8 and 9 (Fig. 3A and B) out of which consistent QTL was detected on chromosome 9 in both the years. The detected linkage of QTL, *qLR*_*9.1*_ controlling leaf rolling character was located on chromosome 9 and observed consistently in both the years in the same physical position on the chromosome under reproductive stage drought tolerance (Fig. 3B; Table 6). The LOD value of 8.21 and phenotypic variance of 65.99 were obtained for the trait from the pooled data analysis (Fig. 3B; Table 6). A distinct peak with additive effect of −1.82 was observed in the graphical representation of QTL analysis of *qLR*_*9*.1_ in the linkage map (Fig. 3B). Position of *qLR*_*9*.1_ was at 22.8cM within the marker interval of RM316 and RM257 on chromosome 9 (Table 6). In case of *qLR*_*8*.1_, the QTL was detected on chromosome 8 (Fig. 3A) within the marker interval of RM337-RM72 and found only in a single experimental year, 2014. The LOD and PVE (%) of *qLR*_*8*.1_ was 2.78 and 60.74, respectively (Table 6).

Similarly, QTL for leaf drying was detected on chromosome 9 and 12 (Fig. 3B). However, the QTL on chromosome 9 was found to be consistent which appeared linked in both the experiment years on the same location (Table 6). From the analysis of result, *qLD*_*9*.1_ controlling leaf drying character was located on chromosome 9 under reproductive stage drought tolerance for both the years (Fig. 3B). A high significant LOD value of 27.85 and phenotypic variance of 80.18 were obtained for the QTL, *qLD9.1* from the pooled data analysis (Fig. 3B; Table 6). A distinct peak was observed in the graphical representation of QTL analysis for *qLD*_*9*.1_ in the linkage map with additive effect of −1.81 (Fig. 3B). Position of *qLD*_*9*.1_ was at 14.8cM within the marker interval of RM316 and RM257 on chromosome 9 (Table 6). In case of the QTL, *qLD*_*12*.1_ which was detected on chromosome 12 (Fig. 3C) within the marker interval of RM20A-RM511 was found only in a single experimental year I (Wet season, 2014). The LOD and PVE (%) of *qLD*_*12*.1_ were 11.35 and 71.72, respectively (Table 6).

The QTL, *qHI*_*9*.1_ located on chromosome 9 at 11.8cM position within the marker interval of RM316 and RM257 was detected to be linked to the trait harvest index under drought stress at reproductive stage of rice (Fig. 3B; Table 6). Additive effect obtained from the study was 0.01 which was minimal effect in this QTL mapping (Fig. 3B). The linkage was detected with a LOD value of 5.21 showing phenotypic variance % of 56.45 for the trait, harvest index (Fig. 3B). Analysis using the software detected the QTL linked to the trait in both experiment years (2014 and 2015). The location on chromosome 9 was same in both the years’ phenotypic data for *qHI*_*9*.1_ (Table 6). In addition, *qSF*_*9*.1_ controlling spikelet fertility was detected as an effective QTL showing LOD value of 3.58 with PVE% of 54.5 (Fig. 3B; Table 6). A clear peak was found in the linkage group at 23.8cM position within the maker interval of RM316-RM257 (Fig. 3B). QTL for relative water content (*qRWC*_*9.1*_) was found at the position of 18.8cM on the basis of wet season 2014 RWC phenotyping data. A significant peak with LOD of 4.13 and PVE% of 59.62 were observed in the marker interval of RM316 and RM257 (Fig. 3B; Table 6). On the basis of wet season, 2015 RWC data, the QTL (*qRWC*_*9.1*_) was again detected within the same marker interval but at a distance of 23.8cM on the chromosome. The LOD value and PVE (%) were found to be 4.78 and 62.65%, respectively in the second year (Fig. 3B; Table 6). However, from the pooled data analysis, it was estimated to be present on chromosome 9 at 21.8cM with LOD value and PVE (%) of 4.27 and 60.87, respectively within the same marker interval. The additive effect of the QTL was found to be 14.56.

## Discussion

The uniqueness of this study was to identify QTL(s) responsible for drought tolerance during reproductive stage. The present experimental design revealed important information on the morpho-physiological basis of different drought related traits under two years, year I (wet season, 2014) and year II (wet season, 2015). Morpho-physiological parameters of 190 RILs and parents under reproductive stage drought stress showed significant variations. Also, the parameters showed significant correlations values among themselves under the stress. A continuous frequency distribution curve for plant height, panicle length, harvest index, grain yield and cell membrane stability was observed from the zymogram (Fig. 1).

Out of the eleven physiological traits studied, five traits *viz*., leaf rolling (*qLR*_*9.1*_), leaf drying (*qLD*_*9.1*_), harvest index (*qHI*_*9.1*_), spikelet fertility (*qSF*_*9.1*_) and relative water content (*qRWC*_*9.1*_ were tagged by QTL composite interval mapping. Chromosome 9 is found to be the most important chromosome showing linkage with five different morpho-physiological traits. Basing on the phenotyping data of year I (wet season, 2014), QTLs namely *qLR*_*9.1*_, *qLD*_*9.1*_, *qHI*_*9.1*_, *qSF*_*9.1*_ and *qRWC*_*9.1*_ were observed in the marker interval of RM316 and RM257 at centimorgan position of 25.8, 13.8, 12.8, 35.8 and 18.8, respectively on chromosome 9. LOD values for the phenotypic traits *viz*., LR, LD, HI, SF and RWC were 7.20, 17.18, 3.51, 2.26 and 4.13, respectively. These QTLs were found to be consistent over both the years.

In field experiment, delaying in leaf drying and leaf rolling are the necessary characteristics for drought tolerance mechanism. Under extreme stress condition, scores for leaf drying generated at all growth stages of rice for determining drought tolerance need to be in moderate to high score. Hence, these consistent QTLs should be pyramided in the recurrent parent for improvement of drought tolerance at reproductive stage in rice. Previous mapping results of leaf rolling under drought by [23] had suggested four QTLs for drought sensitivity index detected, *lr*_*8.1*_, *lr*_*4.1*_, *lr*_*10.1*_, and *lr*_*12.1*_. Also, earlier finding suggests that QTL (*lr*_*8.1*_) for relative water content and leaf rolling was already mapped at equivalent regions (near RM72) on chromosome 8 [18]. Experimental findings of [25] suggested that, under drought stress, 3 markers (RM212, RM302 and RM3825) linked to morphological trait, leaf rolling on chromosome 1. These three markers were also linked to leaf drying under drought stress. As per findings of [21], 5 QTLs were detected for leaf rolling trait out of which one QTL was in chromosome 9 at a distance of 65.6cM. Also in another experiment conducted under drought stress condition, phenotypic variation of 24.8% detected for the leaf rolling QTL on chromosome 6 [27]. Under drought stress situation, chromosome 12 contains another QTL for leaf rolling trait near RM101. However, in this study mapped QTL for leaf rolling were located on chromosome 9 at a position of 22.8cM with LOD of 8.21, PVE(%) of 65.99 and additive effect of −1.82 and hence a novel QTL controlling reproductive stage drought tolerance in rice. Our mapping results detected another QTL for leaf rolling trait, located in the marker interval of RM72 and RM337 at a distance of 27.1cM (Fig 3B) on chromosome 8. This QTL has already been reported by [23] and [18]. Hence, this QTL is validated and can be useful for marker-assisted breeding.

Previous mapping results indicate that RM8085 at 139.9cM linked to leaf drying and leaf rolling trait on chromosome 1 under severe drought stress condition [26]. Published results of [21] detected four QTLs for leaf drying one each in chromosome 1, 3, 3 and 11, respectively. Mapping results of [24] also suggests that *qld*_*1.1*_ located on chromosome 1 controls the leaf drying under drought stress. Thus, the current finding of leaf drying in our experiment may be a new QTL found within the marker interval of RM316-RM257, on chromosome 9 with LOD value 27.85, PVE (%) 80.18 and additive effect of −1.81 controlling leaf drying under drought stress and designated as *qLD*_*9.1*_. Our mapping results showed that another leaf drying linked QTL, *qLD*_*12.1*_ is located in the boundary interval of RM20A and RM511 at a distance of 45cM (Fig. 3) on chromosome 12. As no QTL is reported in this region of the chromosome, the detected QTL, *qLD*_*12.1*_ controlling leaf drying under drought stress with a percentage of phenotypic variance 71.72 may be a novel QTL identified.

Consistent QTLs for harvest index, straw yield, and grain yield under drought stress were reported earlier for RM314 on chromosome 6 with the experiments regulated for several seasons [27]. Earlier study on mapping suggested that in stress situation RM315 is associated with harvest index trait on chromosome 1 [17]. A QTL designated as *qhi*_*1.1*_, expressed only under extreme drought situation was reported on chromosome 1 [20]. A QTL with 65.6% phenotypic variance for harvest index under stress was detected on chromosome 2 [24]. As per their reports, the QTLs for epicuticular wax, harvest index, and water loss rate from processed cut leaves under stress were detected to be co-located with QTLs related to root and shoot related drought tolerance characters in particular rice progenies and might be functional for rain-fed rice development. As under drought situation, the correlation among grain yield (GY) and harvest index (HI) showed high value specifying that HI is a principal element of grain yield under stress. Therefore, genetic advancement of HI would require for enhancement of GY [36, 37]. The QTLs on chromosome 3 classified as *qhi*_*3.4*_, *qhi*_*3.2*_, *qhi*_*3.1*_, *qhi*_*3.3*_, and *qhi*_*3.5*_ were reported with the marker interval of RG104-RM231 position [20]. Another QTL reported on chromosome 1 specified as *qhi*_*1.1*_ that expresses only under severe stress condition [20]. Therefore, the QTL detected in our experiment for controlling HI and located on chromosome 9 within the marker interval of RM316-RM267, LOD of 5.21, PVE(%) of 56.45 and additive effct of 0.01 under drought stress may be a new QTL and designated as *qHI*_*9.1*_.

Grain yield showed positive correlations with spikelet fertility under drought stress at reproductive stage. When spikelet fertility decreased, grain yield also consequently decreased. Pollen formation in rice plants is highly sensitive to drought stress. Abiotic stress during meiotic stage results pollination failure, pollen sterility leading to zygotic abortion and finally spikelet death, but female fertility is affected only under extreme stress. Previous publication of three QTLs, designated as *qpss*_*5.1*_, *qpss*_*4.1*_, and *qpss*_*9.1*_ reported on chromosomes 5, 4, and 9 were under the irrigated situations [20]. Under slight water stress condition, the QTLs *qpss*_*5.3*_ and *qpss*_*5.2*_ were detected on chromosome 5. The QTLs responsible for *qpss*_*5.3*_ and *qpss*_*5.2*_ were detected in the same marker interval of EM15_4 to CDO20, as *qpss*_*5.1*_. Under more water stress level, the QTLs like *qpss*_*8.2*_ and *qpss*_*8.1*_ were detected. The *qpss*_*8.2*_ and *qpss*_*8.1*_ were both detected in the marker interval of ME5_4 to RM256. The QTL detected by us, *q*SF_9.1_ under water stress is nearer to the location of *qpss*_*9.1*_ reported by [20]. The peak marker RM219 for *qpss*_*9.1*_ was in 11.7cM position which is not far from our QTL *qSF*_*9.1*_ located at 12.8cM position in the same chromosome. A QTL was reported on chromosome 9 for spikelet sterility [38]. However, this quantitative trait locus was located at different position of the reported QTL, *qpss*_*9.1*_. The two QTLs, *qpss*_*8.2*_ and *qpss*_*8.1*_, which showed its effect under moderate and extreme drought stress, were also linked with OA (*oa*_*8.1*_) QTL at the same position as per report of [38]․. While [19] detected QTL for OA, at RG1-RM80 marker interval. Mapping results of [13] showed four QTLs namely *qRsf3*, *qRsf5*, *qRsf8* and *qRsf9* controlling spikelet fertility under drought conditions. Thus the result concluded that our QTL, *q*SF_9.1_ might be present in the same locus on chromosome 9 related to spikelet fertility which can further used for the validation of different breeding approaches.

Under water stress condition, higher value of relative water content (RWC) is obtained from drought tolerant genotypes at reproductive stage as compared to susceptible genotypes. Earlier mapping work of [36] indicated that the marker interval R2417-RM212-C813 on chromosome 1 was associated with RWC under field drought condition. A QTL, *qrwc*_*11.1*_ in chromosome 11 in between marker interval of RM254-RG1465 reported for controlling the RWC in rice [24]. [3] reported the physiological response of relative water content affecting grain yield in *qtl*_*12.1*_ under drought condition. Mapping population developed for map physiological trait including relative water content by [18] map RWC trait on chromosome 1, 3 and 8. Another linkage map by [39] constructed in chromosome 1, 6 and 12 form marker interval of RG810-RG348 for relative water content trait. As RWC trait is showing its consistent effect for the both the years (wet season, 2014 and wet season, 2015) on chromosome 9 in between the marker interval of RM316-RM257, 21.8cM position and this trait was not map before, thus *qRWC*_9.1_ can be consider as a novel QTL from our experiment. Experimental result conforms that the LOD value of 4.27 and PVE (%) of 60.87 and additive effect of 14.56 (Table 6) was obtained from the pooled data analysis result. Therefore, the QTL detected in this experiment for controlling RWC under drought stress may be a novel QTL and designated as *qRWC*_*9.1*_.

We observed a significant correlation of plant height (r=0.209**), panicle length (r=0.252**), panicle emergence (r=0.201**), thousand seed weight (r=0.302**) with grain yield. These traits were reported earlier as important drought controlling factors responsible for growth, development and yield parameters in rice [40]. However, there were no QTL found linked to these traits from our mapping study. Cell membrane stability (CMS) is considered to be one of the major selection indices of drought tolerance in cereals [16]. A negative correlation of −0.134 for CMS was observed for grain yield from this study. But, no QTL was detected for this trait.

## Conclusion

The present study showed the nature of correlation of physiological parameters with themselves and with grain yields under reproductive stage drought stress. Five consistent QTLs were detected *viz., qLR*_*9*.1_, *qLD*_*9*.1_, *qHI*_*9*.1_, *qSF*_*9*.1_, and *qRWC*_*9*.1_ controlling leaf rolling, leaf drying, harvest index, spikelet fertility and relative water content, respectively during reproductive stage drought stress. Also, *qLR8.1* was validated in this mapping population and useful in marker-assisted breeding (MAB) programs. The correlated physiological traits may be useful in selecting desirable drought tolerant plants for reproductive stage drought stress. Also these identified QTL regions can be useful in marker-assisted breeding program for development of drought tolerant rice plant.

## Acknowledgments

The authors are highly grateful to the Director, ICAR-National Rice Research Institute, and Head, Crop Improvement Division of the Institute for providing all the necessary facilities.

## Author Contributions

Conceptualization: SKP

Data curation: SRB.

Formal analysis: SRB

Funding acquisition:SKP.

Investigation: SRB, EP, SPM.

Methodology: SKP SRB.

Project administration: SKP.

Resources: SKP

Software: SRB

Supervision: SKP TM

Writing - original draft: SKP SRB.

Writing - review & editing: SKP TM

